# The design and evaluation of a Bayesian system for detecting and characterizing outbreaks of influenza

**DOI:** 10.1101/435727

**Authors:** Nicholas E. Millett, John M. Aronis, Michael M. Wagner, Fuchiang Tsui, Ye Ye, Jeffrey P. Ferraro, Peter J. Haug, Per H. Gesteland, Gregory F. Cooper

## Abstract

The prediction and characterization of outbreaks of infectious diseases such as influenza remains an open and important problem. This paper describes a framework for detecting and characterizing outbreaks of influenza and the results of testing it on data from ten outbreaks collected from two locations over five years. We model outbreaks with compartment models and explicitly model non-influenza influenza-like illnesses.

## 2 Introduction

The prediction and characterization of outbreaks of infectious diseases remains an open and important problem [1]. Influenza, with nearly annual outbreaks in temperate regions of the world, provides an ideal test domain [2].

This paper describes a framework for detecting and characterizing outbreaks of influenza and the results of testing it on data from ten real outbreaks collected from two locations over five years. Like several other systems, we model outbreaks with compartment models [3, 4, 5]. We differ from this past work in that we use the full text of patient care reports, rather than just chief complaints [6], counts of syndromes from sentinel physicians [5], counts of internet queries [7], etc. Doing so provides a rich source of evidence that may provide an early signal of an outbreak.

We use the evidence to reason about likelihoods, such as *P*(*findings*|*influenza*) or *P*(*findings*|*RSV*), rather than just simple counts. The approach is quite general, since the findings can include any evidence about a patient’s disease status, including history, symptoms, signs, labs, and other information.

This paper extends our previous work [8] by using a more sophisticated model of non-influenza influenza-like illness (NI-ILI), modeling a probability distribution over influenza outbreak start dates, and testing on a set of real outbreak data collected over five years at two locations widely separated in the United States.

## 3 System Architecture

We have developed an end-to-end framework for outbreak detection and characterization [9]. It starts with patient care reports, extracts findings with natural language processing (NLP), assigns likelihoods to each patient case with a case-detection system (CDS), and constructs a model with an outbreak-detection system (ODS) that can be used for prediction and characterization.

A patient’s care report contains the most detailed and complete record of their present illness available. Much of the information in it (including chief complaint, history of present illness, a detailed patient assessment, treatment, and response to treatment) is in free-text. Other information, such as laboratory findings, is codified. In our system, such data, including symptoms and signs, are extracted using natural language processing software [10]. Some patient care reports include a laboratory test for influenza which can provide a definitive diagnosis of influenza.

The findings (free-text derived and coded) for each patient are passed to CDS which derives the probability of those findings given each of *influenza*, *NI-ILI*, and *other*. *NI-ILI* implicitly includes several diseases, such as respiratory syncytial virus (RSV) and parainfluenza, and *other* includes everything else such as trauma, appendicitis, etc. CDS uses a Bayesian network that represents the joint probability distribution of each patient’s findings (including laboratory results) and the three disease categories just mentioned [11]. As mentioned, it provides the likelihoods *P*(*E*(*p*, *d*)|*influenza*), *P* (*E*(*p, d*)|*NI-ILI*), and *P* (*E*(*p, d*)|*other*), where *E*(*p, d*) is the set of findings for patient *p* on day *d*. That is, the probability of the patient’s findings given they have each of *influenza*, *NI-ILI*, or *other*.

ODS takes all of the evidence from the first day of the monitored period through the present, evaluates thousands of possible outbreak models against the data, and makes projections about the future. Let 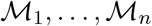 be a representative set of models and *E*(1: *c*) be all of the data available through the current day *c*. ODS computes the expected number of *influenza* cases on each day *d*, with:

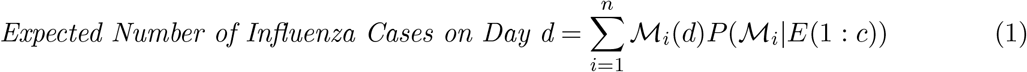

where 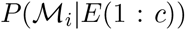 is the probability of model 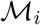 given the data up to the current day, and 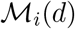 is the number of *influenza* cases predicted by model 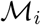 on day *d*. Typically, *d > c* since we want to predict the future, but we can assess the past with *d < c*. ODS is based on an earlier system described in [8].

Most patients spend only a few hours in the ED and their report is completed and available shortly after they leave. However, a significant number of reports take several days to complete (sometimes spread over multiple follow-up reports) and are not immediately available to a real-time system. We avoid *data leakage* (when the system is tested using data that are theoretically—but not actually—available each day [12]) by using only the records that are available at the time of evaluation, then retroactively augmenting previous evidence as records become available, and recomputing previous likelihoods.

## 4 Modeling Influenza

We model immunizing infectious diseases like *influenza* with SEIR models [13] that divide the population into four compartments:

*Susceptible:* Individuals who could become infected.
*Exposed:* Individuals who have the disease but cannot yet transmit it.
*Infectious:* Individuals who have the disease and can transmit it.
*Recovered:* Individuals who are immune to contracting the disease.

Typically, a large part of the population is initially *susceptible*, a significant number may be *recovered*, and the infection is introduced by a small number of *infectious* individuals.

After the entire population has been partitioned into these four compartments, they move through the compartments according to several parameters:

*R*_0_ is the expected number of individuals that an infectious person would infect if he were introduced into an entirely susceptible population.
*Latent Period* is the average amount of time before an exposed individual becomes infectious.
*Infectious Period* is the average amount of time an individual is infectious.

When modeling an outbreak of *influenza*, ODS generates thousands of possible SEIR models and uses each one according to Equation 1. These models are generated by randomly and uniformly drawing values from Table 1. These ranges have been determined by expert opinion and past outbreaks of as reported in the literature *influenza* [13, 2, 14].

**Table 1:** SEIR Parameter Ranges

| Parameter | Minimum Value | Maximum Value |
| --- | --- | --- |
| *Initial S* | $0.0 * population$ | $1.0 * population$ |
| *Initial E* | 0 | 0 |
| *Initial I* | 1 | 100 |
| $R_0$ | 1.1 | 1.9 |
| *Latent Period* | 1 | 3 |
| *Infectious Period* | 1 | 8 |

Although ODS models *influenza* in the general population, its evidence comes from hospital ED’s, which represent only a fraction of the actual number of *influenza* cases. To adjust for that difference, we estimate that 1% of people infected with *influenza* in the general population actually go to an ED based on estimates reported in [15].

## 5 Outbreak Detection and Characterization

Equation 1 relies on the value 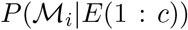 for each 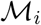. We can find this probability using Bayes’ rule:

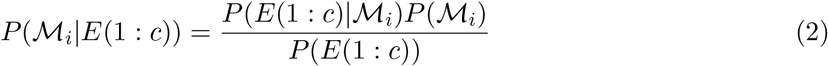

We can interpret 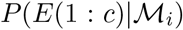 as how well 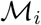 explains the data. Note that *P*(*E*(1: *c*)) is the same for every model and 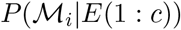 is proportional to the numerator.

### 5.1 The Prior Probability of a Model

In temperate climates, such as the United States, outbreaks of *influenza* occur nearly every year. If an outbreak does occur, it can peak as early as October 1 or as late as April 1, but typically *influenza* outbreaks peak during the winter months.

Peak dates depend on start dates. We model the distribution of start dates with the following procedure:

1. Create 100, 000 models with start dates between June 1 and the following March 1.
2. Save the set *S* of models with peak infectious counts that occur between October 1 and April 1 of the following year.
3. Create a probability distribution over the start dates in *S* by setting *P*(*d*) to the fraction of models in *S* that start on day *d*.

We assume that the probability an influenza outbreak will occur in any given year is 0.9. We create a special non-outbreak model, 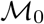 with 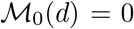 for any day *d*, and set 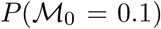. Let 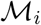 be an outbreak model that starts on day *d*. For each possible start day *d*, ODS samples and evaluates *n* equiprobable outbreak models that start on that day, so:

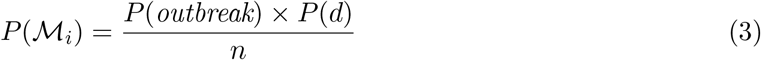

where *P* (*outbreak*) = 0.9.

### 5.2 The Posterior Probability of a Model

In Equation 2 we need 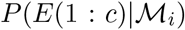 for each model. Assuming that the evidence on each day *E*(*d*) is independent on any other day given a model of *influenza*, we have:

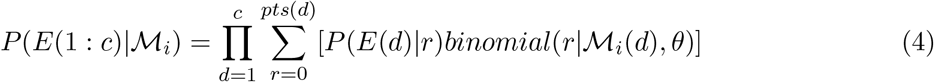

Here, *pts*(*d*) is the total number of patients in the ED on day *d*, 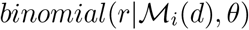 is the probability that exactly *r* people with *influenza* go to the ED given there are 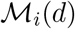 people in the population who independently choose to go the ED with probability *θ*. We sum over all the possibilities.

If we assume that on each day the patients in the ED present independently of each other, we have:

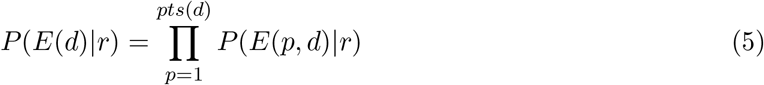

That is, the probability of all of the evidence *E*(*d*) on day *d* is the product of the probability of the evidence of each patient *E*(*p, d*) on that day.

Finally, we need the probability of each patient’s evidence given there are *r influenza* patients in the ED. If we assume that *influenza*, *NI-ILI*, and *other* are mutually exclusive and exhaustive for a single patient, then:

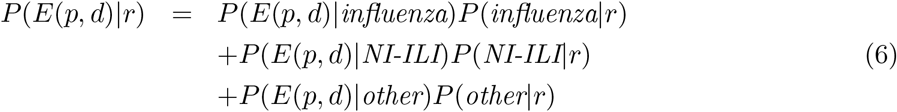

We estimate *P* (*influenza*|*r*) as *r/pts*(*d*). We describe our method to estimate *P*(*NI-ILI*|*r*) later. Finally, *P*(*other*|*r*) = 1 − (*P*(*influenza*|*r*) + *P*(*NI-ILI*|*r*)).

### 5.3 The Probability that an Outbreak is Occurring

The probability that an outbreak has started by day *c* is simply 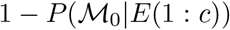 where 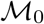 is the non-outbreak model. (Recall that 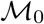 is the non-outbreak model with 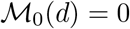 for each day *d*.) But even if an outbreak has started by day *c*, that does not mean that it is occurring on day *c*, since it may have already ended. We want the probability that some outbreak is occurring on day *c*.

The probability that an arbitrary individual in the population has *influenza* on day *c* given model 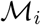 is:

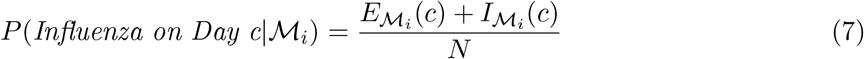

where 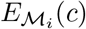 is the number of exposed people and 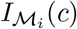 is the number of infectious people on day *c* for model 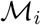, and *N* is the total population in the region being monitored. We can now compute the probability that an outbreak is occurring on day *c* for model 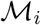 as:

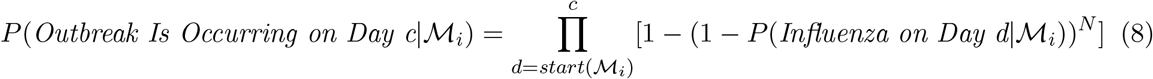

The term 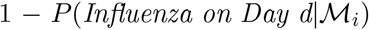 is the probability that an arbitrary individual in the population is not infected, 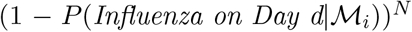 is the probability that nobody is infected, and 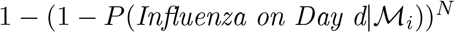 is the probability that at least one individual is infected, given model 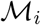. The product is the probability that at least one individual in the population is infected on each day from the start of 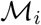 to the current day *c*. Finally, the probability there is an ongoing outbreak on day *c* is:

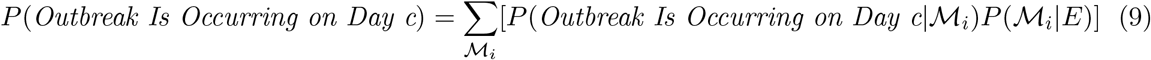

where 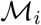 ranges over the entire set of models.

### 5.4 Estimating NI-ILI

By definition, *NI-ILI*—*non-influenza influenza-like illness*—presents like *influenza*. Furthermore, some forms of *NI-ILI*, such as RSV, display outbreak activity. Properly recognizing outbreaks of *influenza* requires us to account for *NI-ILI*. (Equation 6 specifies how we incorporate an estimate of *NI-ILI* to score models.) Appendix A describes a technical approach to estimating the level of *NI-ILI* during the summer and throughout the *influenza* season.

## 6 Experimental Results

Our datasets consisted of 5 years of data from 2010-2015 for both Allegheny County (AC), Pennsylvania and Salt Lake County (SLC), Utah. We assume that a new *influenza* season starts on June 1 of a given year. Thus, the first season is defined to span from June 1, 2010 through May 31, 2011, the second season spans from June 1, 2011 through May 31, 2012, and so on through May 31, 2015.

Our gold standard for determining the actual peak of an outbreak in the population is laboratory-confirmed cases of influenza associated with emergency department visits. According to this standard, the peak of an outbreak is the maximum of a seven day central moving average of the daily positive lab test counts. We refer to this as the *laboratory-peak*.

Appendix B shows the counts of positive *influenza* lab tests, along with the CDS expected number of ED *influenza* cases, for each *influenza* season from 2010-2015 for both regions. These results give an indication of the relative size and shape of each outbreak. Note that these include graphs that appear symmetric and look like SEIR curves (as in Figure 1), graphs that include two distinct peaks (as in Figure 2), graphs that are asymmetric (as in Figure 6), and graphs that display little or no outbreak activity (as in Figure 3).

**Figure 1:**
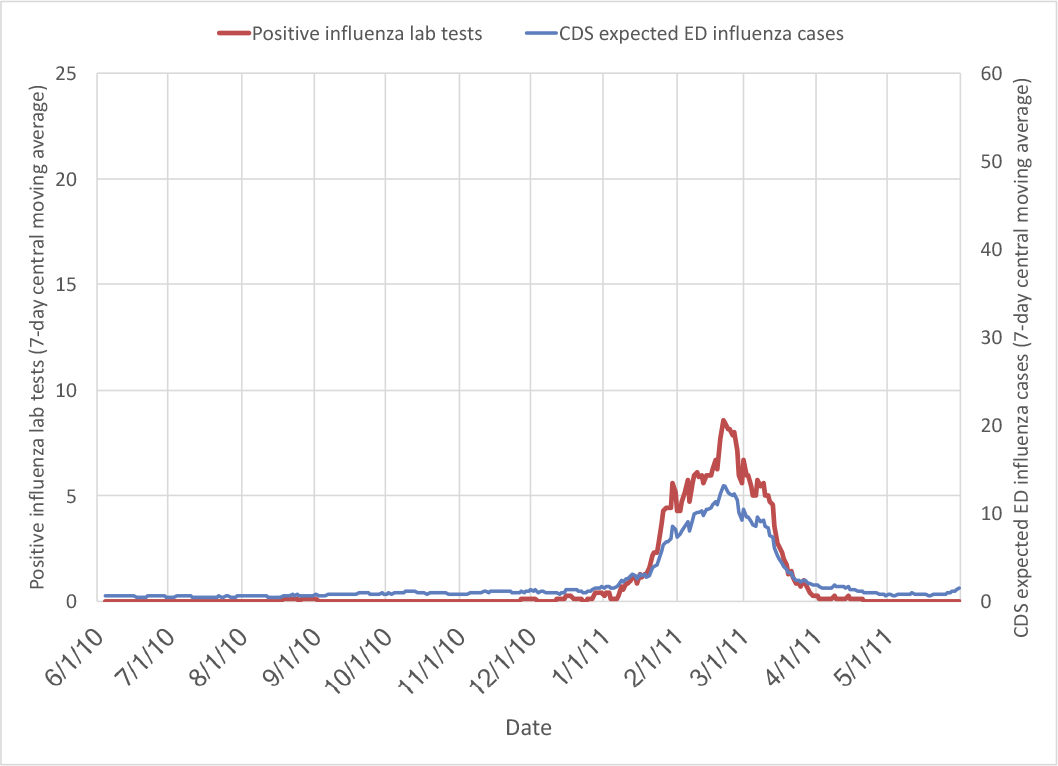
Allegheny County, 2010-2011

**Figure 2:**
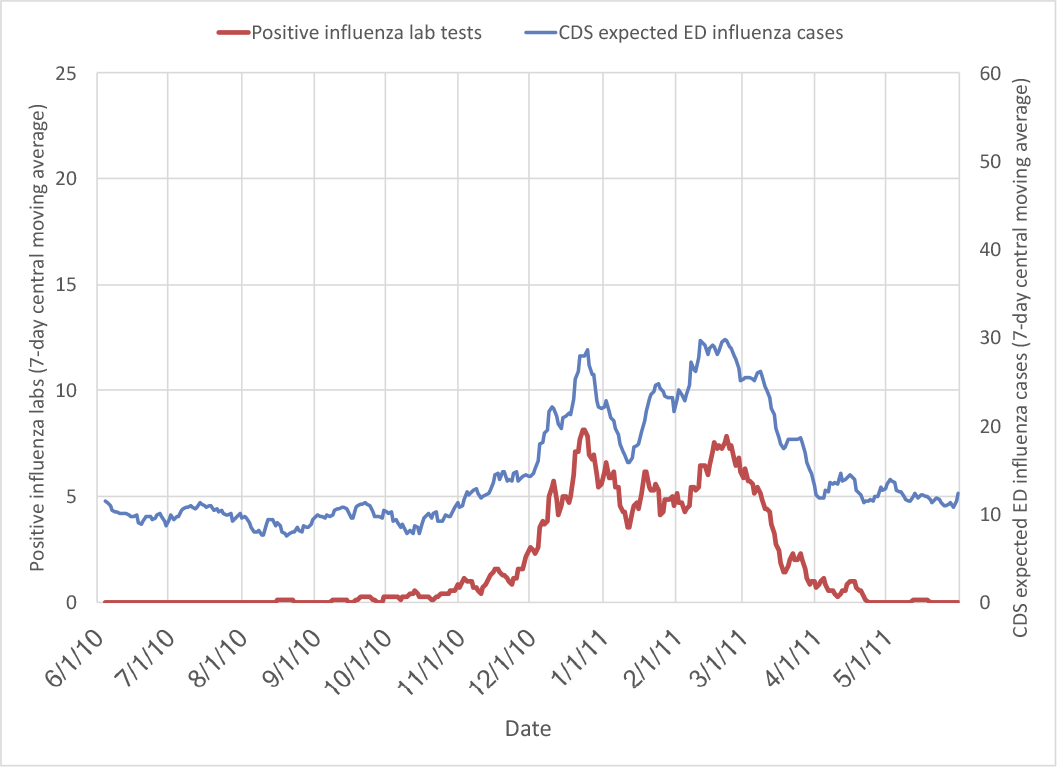
Salt Lake County, 2010-2011

**Figure 3:**
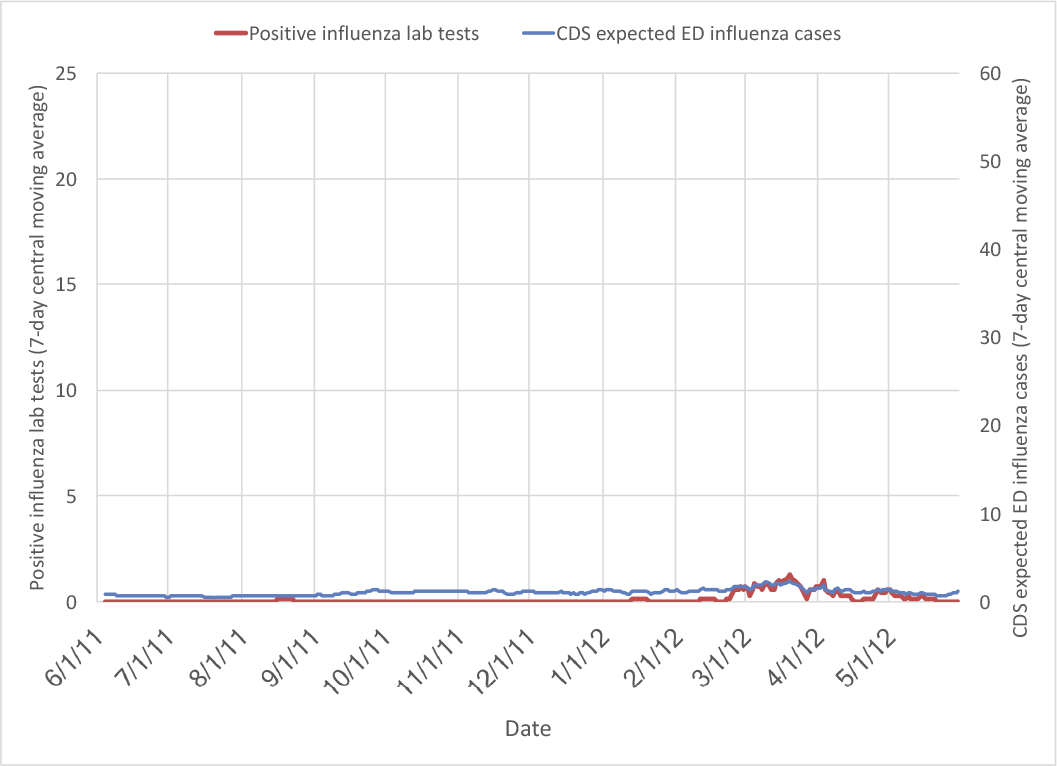
Allegheny County, 2011-2012

**Figure 4:**
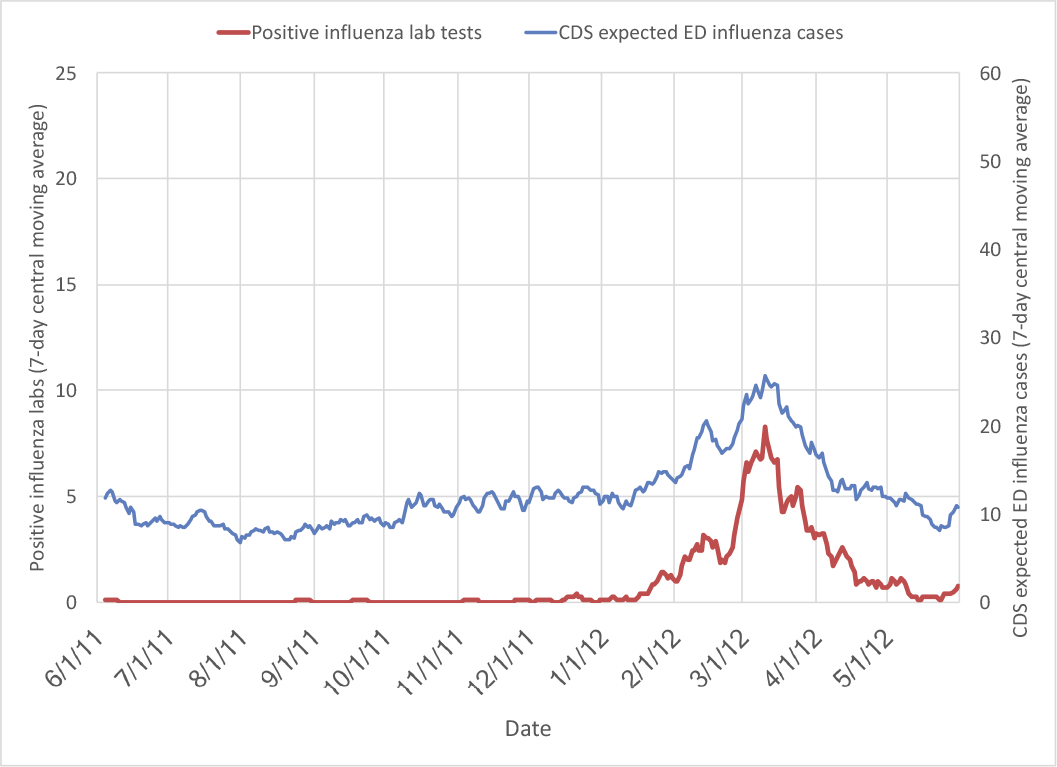
Salt Lake County, 2011-2012

We ran ODS on each day from August 2 through the following June 1 for each *influenza* season in each region. On a given day, ODS uses CDS data through the previous day, so running ODS on June 1 uses data through May 31, consistent with our definition of the end of an *influenza* season. We started running ODS on August 2 (using data through August 1) because the data from June 1 through July 31 is used to compute the initial *influenza* and *NI-ILI* priors used by ODS (See Appendix A.)

Our metric for quantifying ODS performance is the absolute error between the ODS-expected peak date and the actual peak date determined from the gold standard (positive lab tests). Table 2 shows the peak date error for each *influenza* season and region at different time points relative to the actual peak date. In parentheses next to each peak date error is the probability of outbreak computed by ODS. In general, we would only trust a peak prediction by ODS when its probability of an outbreak is high at that point. For example, for the 2013-2014 season for AC, at a point 6 weeks before the actual peak, the ODS predicted peak was 19 days after the actual peak. At this point we can see the probability of outbreak was computed to be only 0.29.

**Table 2:** ODS peak date error for each *influenza* season from 2010-2015 for Allegheny (AC) and Salt Lake (SLC) counties. The ODS-computed probability of outbreak is shown in parenthesis next to each peak date error. The error value is the difference between the ODS-computed expected peak date and the actual peak date. Thus, a positive error means that the ODS-predicted peak is later than the actual peak. Predictions made with greater than 50% probability are shown in bold. The average absolute error values are for predictions with greater than 50% probability.

|  |  | Date of prediction relative to actual peak |  |  |  |
| --- | --- | --- | --- | --- | --- |
|  |  | -6 weeks | -4 weeks | -2 weeks | 0 weeks |
| 2010-2011 | AC | **7 (1.00)** | **13 (1.00)** | **7 (1.00)** | **5 (1.00)** |
| 2011-2012 | AC | 18 (0.15) | 25 (0.08) | 33 (0.16) | **43 (0.96)** |
| 2012-2013 | AC | 53 (0.10) | 61 (0.10) | **23 (1.00)** | **28 (1.00)** |
| 2013-2014 | AC | 19 (0.29) | **14 (1.00)** | **19 (1.00)** | **17 (1.00)** |
| 2014-2015 | AC | **8 (0.60)** | **9 (1.00)** | **13 (1.00)** | **8 (1.00)** |
| AC Average Absolute Error |  | 7.5 | 12.0 | 15.5 | 20.2 |
| 2010-2011 | SLC | 14 (0.28) | **2 (1.00)** | **12 (1.00)** | **7 (1.00)** |
| 2011-2012 | SLC | **-4 (0.79)** | **2 (1.00)** | **1 (1.00)** | **6 (1.00)** |
| 2012-2013 | SLC | 41 (0.12) | **-7 (0.74)** | **3 (1.00)** | **7 (1.00)** |
| 2013-2014 | SLC | 48 (0.21) | **3 (1.00)** | **5 (1.00)** | **1 (1.00)** |
| 2014-2015 | SLC | 55 (0.12) | 56 (0.20) | **5 (1.00)** | **-1 (1.00)** |
| SLC Average Absolute Error |  | 4.0 | 3.5 | 5.2 | 4.4 |
| Overall Average Absolute Error |  | 6.33 | 7.14 | 9.78 | 12.3 |

ODS sampled 100 models per day on each of the 274 possible outbreak start days for a total of 27,400 outbreak models and one non-outbreak model. ODS required approximately 45 minutes of runtime to analyze all the data for a given year (corresponding to a row in Table 2) when using a Macbook Pro with a single Intel 2.2GHz i7 processor.

The research protocol was approved by both institutional IRBs (University of Pittsburgh: PRO08030129; Intermountain Healthcare: 1024664). All patient data were de-identified and analyzed anonymously. No consent was given.

## 7 Limitations

The work reported here has some limitations. The data is limited to hospital emergency departments (EDs) and does not include clinics or private practices. Furthermore, these EDs are urban and suburban, and do not include rural areas. ODS relies on natural language processing software to extract findings from patient care reports and a case-detection system to produce likelihoods, so we rely on the accuracy of those systems.

We used the number of laboratory-confirmed *influenza* cases as a gold standard because we want to recognize and characterize *influenza*, as opposed to influenza-like illnesses in general. However, the total number of laboratory-confirmed cases of *influenza* is relatively small. Furthermore, we assumed that the rate of testing is constant throughout an outbreak, but it is possible that testing policies change as an outbreak develops.

We also assume that an outbreak can be modeled with a single SEIR model with constant parameters. We will address this assumption below.

## 8 Discussion

Table 2 shows that ODS’s peak date predictions are almost always late. An examination of the graphs of counts of positive influenza lab tests in Appendix B provides a clue why: the outbreaks are not symmetric. (Allegheny County 2012-2013 in Figure 5 is a good example.) To explain this asymmetry, we need to reexamine our basic assumption that there is a single outbreak of influenza each year that can be modeled by a SEIR model.

**Figure 5:**
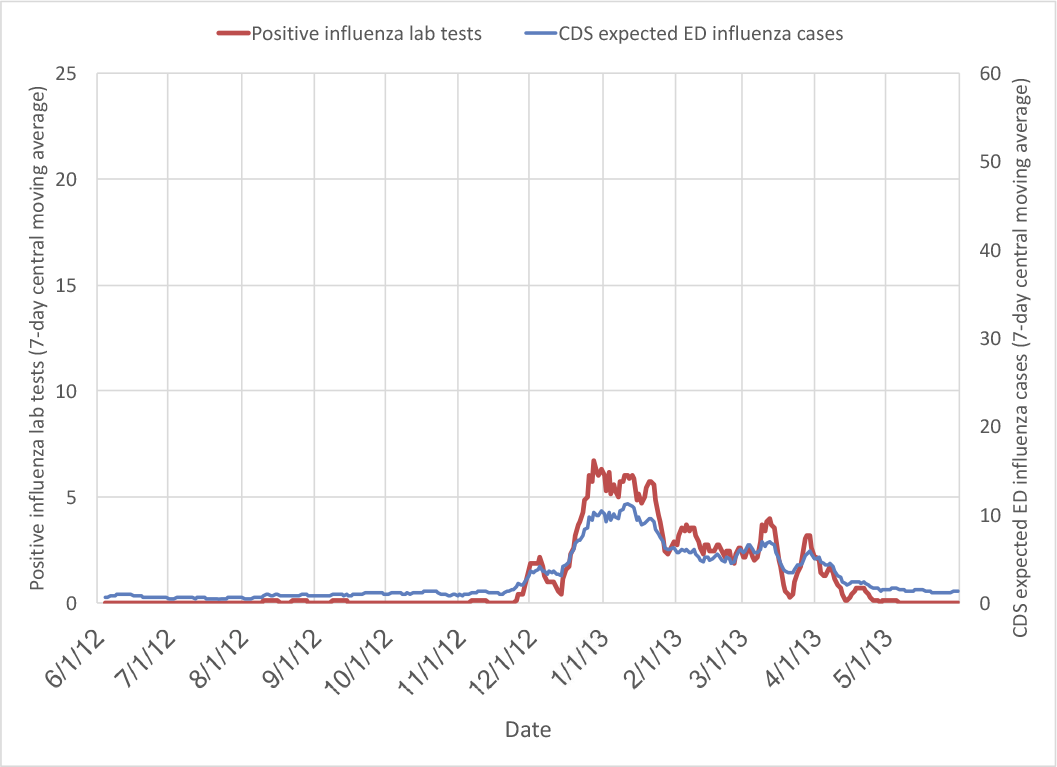
Allegheny County, 2012-2013.

**Figure 6:**
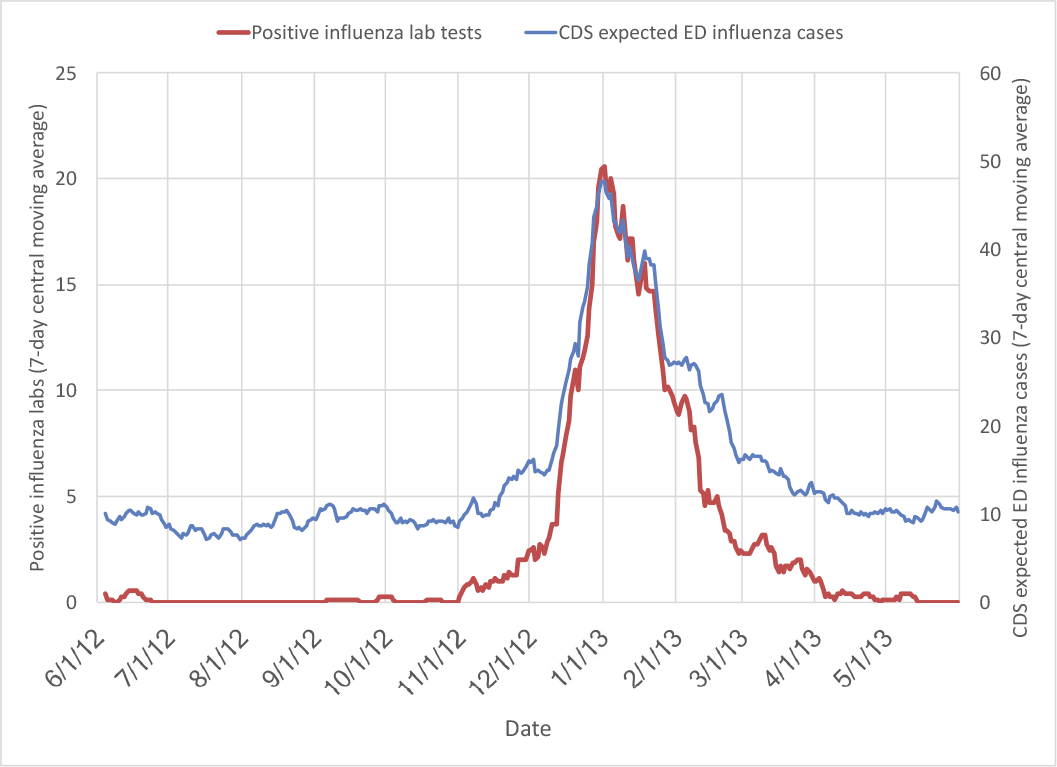
Salt Lake County, 2012-2013

**Figure 7:**
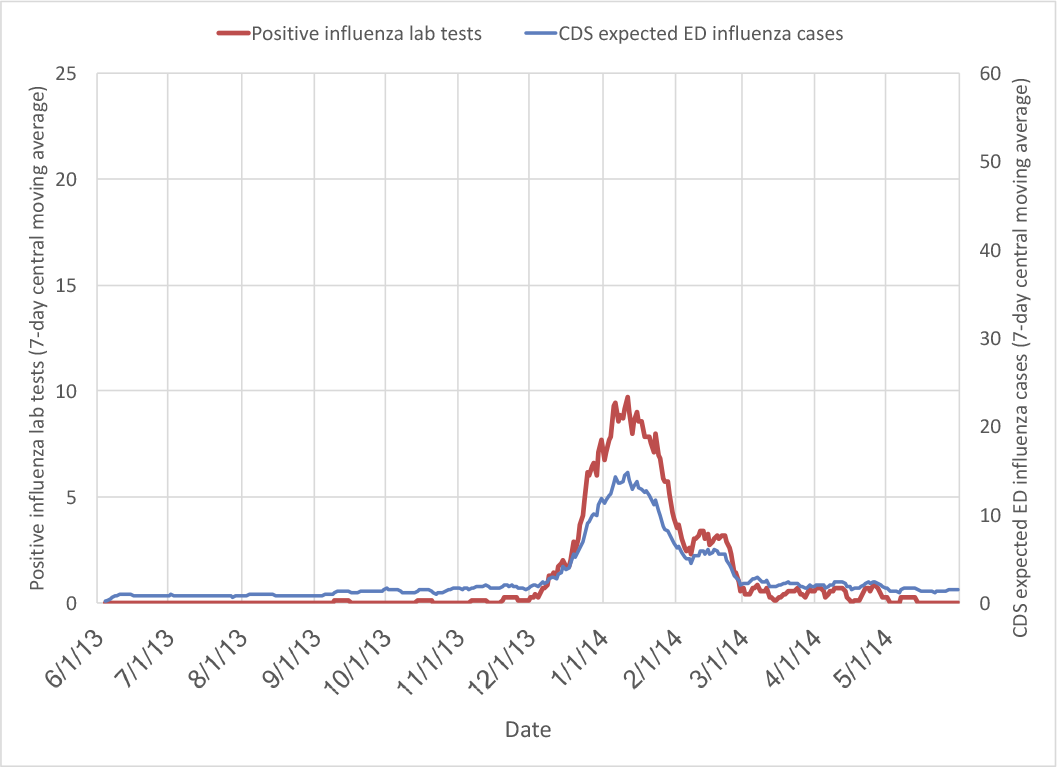
Allegheny County, 2013-2014

**Figure 8:**
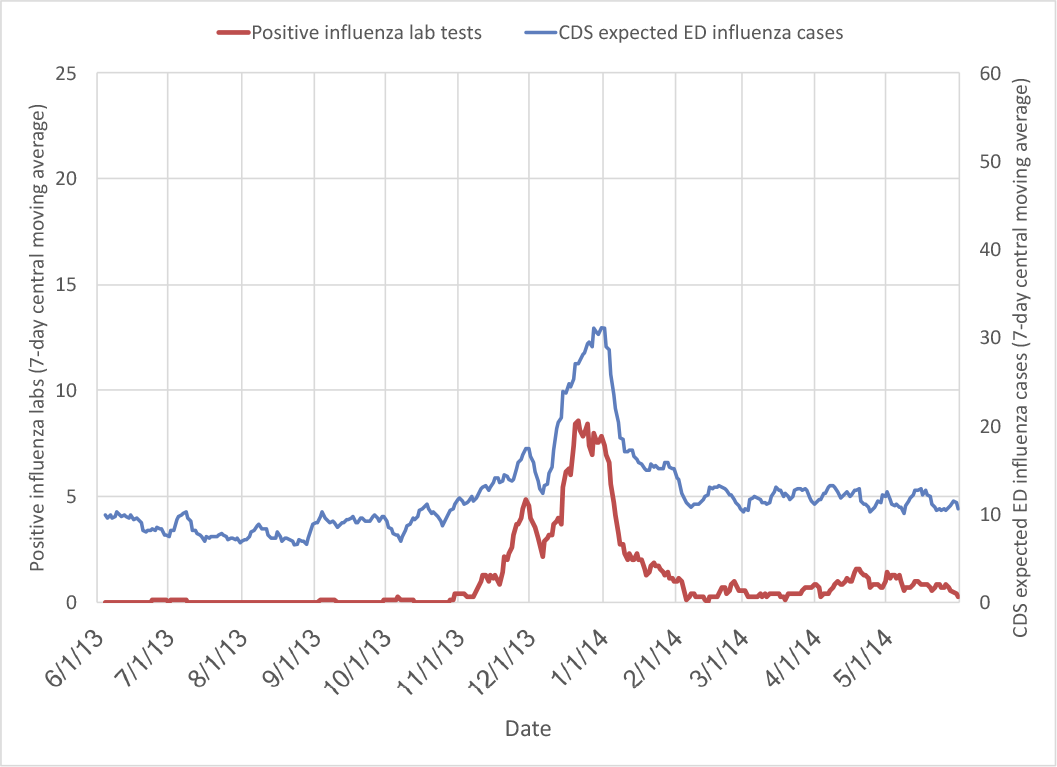
Salt Lake County, 2013-2014

**Figure 9:**
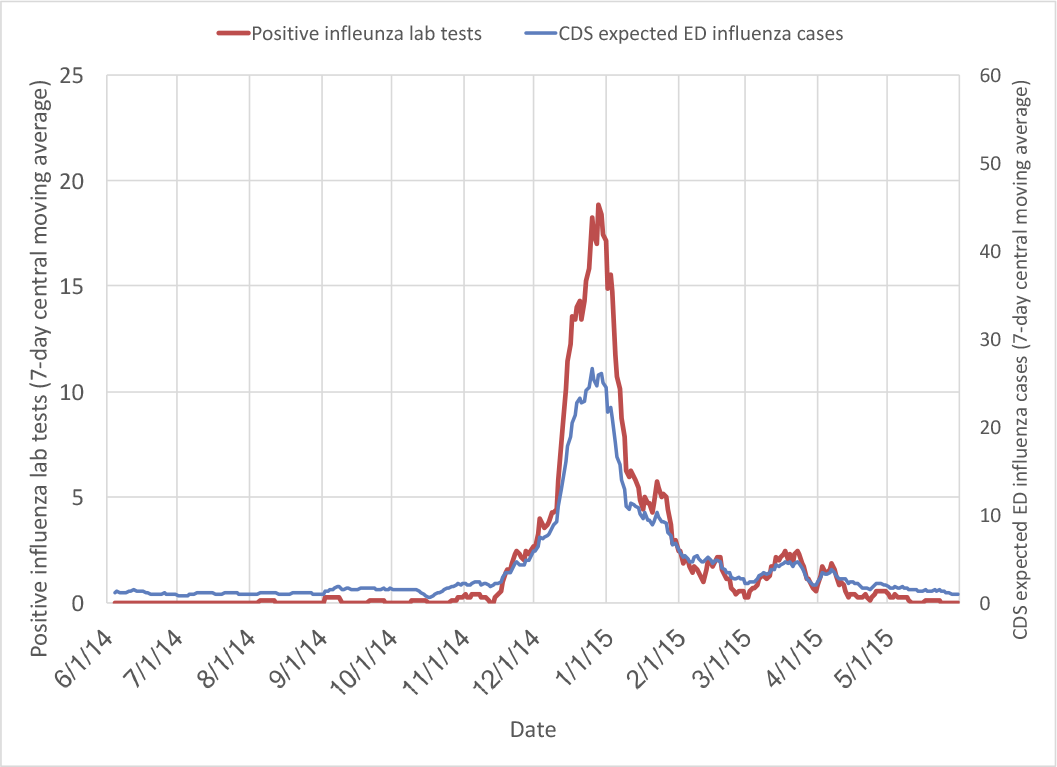
Allegheny County, 2014-2015

**Figure 10:**
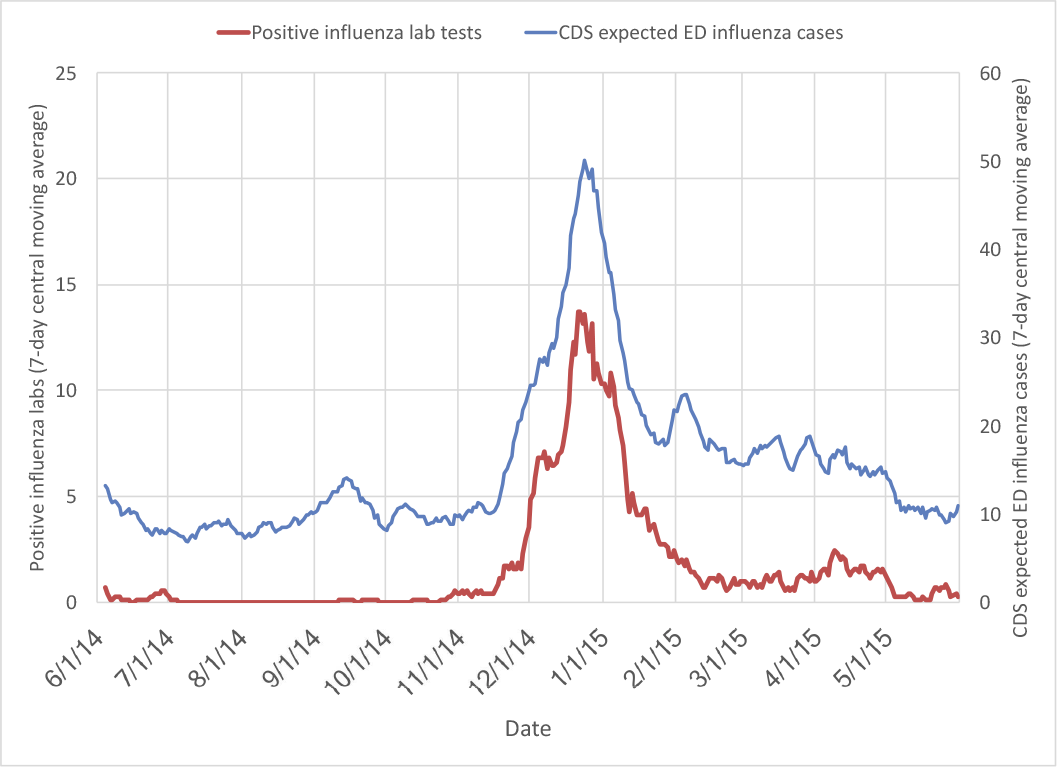
Salt Lake County, 2014-2015

There is rarely a single outbreak of influenza each year. Typically, there is an outbreak of influenza A, followed by a smaller outbreak of influenza B. Table 3 compares the relative sizes of influenza A and B outbreaks for HHS region 3 (which includes Allegheny County) and region 8 (which includes Salt Lake County), according to data from the CDC’s FluView [16]. For instance, the first row states that for the influenza year 2010-2011 in region 3, there were 815 laboratory-confirmed cases of influenza A at its peak, and 148 laboratory-confirmed cases of influenza B at its peak. Also, the influenza B peak was 18% as large as the influenza A peak and occurred one week later. Notice that in Table 2, ODS’s worst results were for Allegheny County 2011-2012, 2012-2013, 2013-2014. Also, in Table 3 *there was a substantial outbreak of influenza B that peaked several weeks after influenza A in exactly those years*.

**Table 3:** Influenza A and B Peaks (see text)

| Year | Region | Influenza A | Influenza B | B/A | Offset (weeks) |
| --- | --- | --- | --- | --- | --- |
| 2010-2011 | 3 (includes AC) | 815 | 148 | 0.18 | 1 |
| 2011-2012 | 3 (includes AC) | 183 | 42 | 0.23 | 5 |
| 2012-2013 | 3 (includes AC) | 1533 | 274 | 0.18 | 11 |
| 2013-2014 | 3 (includes AC) | 566 | 119 | 0.21 | 11 |
| 2014-2015 | 3 (includes AC) | 1757 | 113 | 0.06 | 13 |
| 2010-2011 | 8 (includes SLC) | 391 | 278 | 0.71 | 0 |
| 2011-2012 | 8 (includes SLC) | 419 | 13 | 0.03 | 4 |
| 2012-2013 | 8 (includes SLC) | 780 | 254 | 0.33 | 0 |
| 2013-2014 | 8 (includes SLC) | 976 | 39 | 0.04 | 20 |
| 2014-2015 | 8 (includes SLC) | 1429 | 177 | 0.12 | 14 |

We believe that the presence of a second, smaller outbreak of influenza B explains most of the deviation of ODS’s predictions from the actual peak dates. However, there are other factors that can affect the prediction of the peak of an outbreak. The basic parameters of the model—especially *R*_0_—can change based on social or environmental factors [17]. This could happen, for instance, if individuals in the population self-quarantine or increase hand washing as an outbreak progresses. Immunization programs and individual decisions about whether and when to be vaccinated could also change the course of the outbreak [18]. These changes would have the effect of reducing the number of individuals in the *Susceptible* compartment of the SEIR model. Note that in both of these examples, the original predicted peak is later than the peak when the changes are taken into consideration. To help address this problem, we could learn the relationships between social and environmental conditions and the parameters of a SEIR model. For instance, we could learn the relationship between *R*_0_ and school closings by comparing deviations of the number of infectious from standard SEIR models with known patterns of holidays, school closings, etc. A simple way to learn such a relationship could start with the assumption that the value of *R*_0_ changes daily according to the number of people affected by school closings. That is, *R*_0_(*d*) = *f* (*c*(*d*)), where *c*(*d*) is the number of people affected by school closings on day *d*. It would be reasonable to assume that *f* is a linear function since *R*_0_ = *βDN* where *β* is the rate at which any two individuals come into effective contact, *D* is the duration of infectiousness, and *N* is the population. We can then set the parameters of *f* to the values that maximize *P* (*E*(*d*)|*R*_0_(*d*) = *f*(*c*(*d*))) over all of the available training data. Of course, *R*_0_ depends on more than just social mixing, and *f* may be a nonlinear, multi-parameter function; in that case, we could investigate ways to approximate it.

Properly modeling outbreaks of influenza will require models that incorporate multiple outbreaks as well as parameters that vary according to day-to-day information. The current work provides a baseline from which to proceed, and insights into which directions to take.

## 9 Conclusions

We have described a system—ODS—that can predict and characterize outbreaks of influenza, and the results of testing it on ten datasets collected over five years at two locations. Our analysis showed that ODS can reliably detect the presence of an outbreak, but systematically predicts the peak later than it actually occurred. We believe this is primarily due to the inability of SEIR models to correctly model the influenza A and B outbreaks that occur most years. Further work with ODS, or similar systems, needs to be based on outbreak models that can describe multiple, overlapping outbreaks of influenza.

## 10 Funding

This work was supported by the National Institutes of Health [R01-LM011370] and the National Library of Medicine [T15-LM007059] to John M. Aronis].

## A Modeling NI-ILI

We compute the level of *NI-ILI* in three stages. First, we compute summer baseline levels of *NI-ILI*. Then, we compute the level of *NI-ILI* during influenza season using a Bayesian time-series. Finally, we adjust the level of *NI-ILI* to account for the increased level of apparent *NI-ILI* during an outbreak of *influenza*.

We compute baseline levels of *influenza* and *NI-ILI* in the summer when it is highly unlikely there is an outbreak. Because they present similarly, and one can be explained by the possible presence of the other, we need to compute them in tandem. Let *S* be the set of all possible pairs of *influenza* and *NI-ILI* prior probabilities. Let *s*_*i*_ be a specific pair of priors {*P*_*i*_(*influenza*), *P*_*i*_(*NI-ILI*)}. We have:

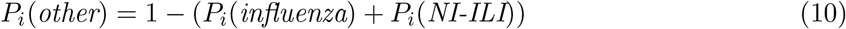

Since the *s*_*i*_ are mutually exclusive and exhaustive, we can compute the probability of the data on day *d* with:

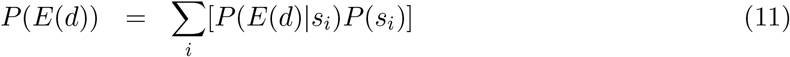

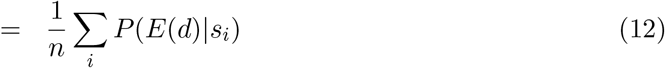

assuming there are *n P*(*s*_*i*_) and they are uniformly equal. We can now compute the expected probability of influenza on day *d* as:

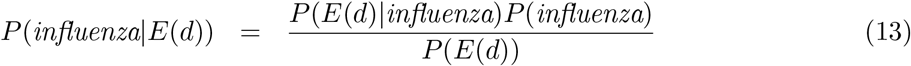

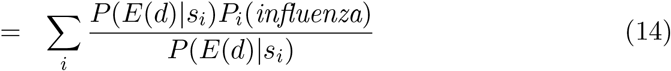

again assuming the *P*(*s*_*i*_) are equal. We can compute *P*(*NI-ILI*|*E*(*d*)) and *P*(*other*|*E*(*d*)) similarly. Finally, let *summer* be all the days of summer:

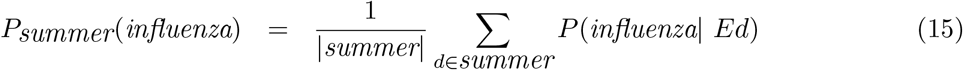

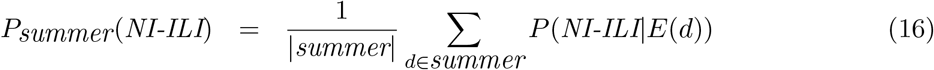

That is, we estimate the background probability of *influenza* and *NI-ILI* as their mean daily probabilities during a summer period.

Given the summer baseline levels of *NI-ILI* and *influenza*, we now compute the probability of *NI-ILI* during the remainder of the year with a Bayesian time-series. Let *E*(*p, d*) be the evidence for patient *p* on day *d*, *E*(*d*) be all of the evidence on day *d*, and *N_d_* be the number of patients on day *d*. Also, let *P*_*d*_(*influenza*), *P*_*d*_(*NI-ILI*), and *P*_*d*_(*other*) denote the prior probabilities for *NI-ILI*, *influenza*, and *other* for day *d*. Let us first assume that we have prior probabilities *P*_*d*_(*NI-ILI*), *P*_*d*_(*influenza*), and *P*_*d*_(*other*) for day *d*. We can then derive the posterior probability of *NI-ILI* for patient *p* on day *d* as:

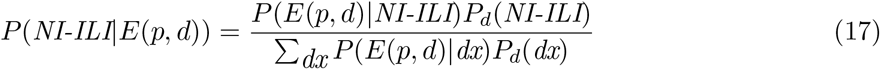

where *dx* ranges over *influenza*, *NI-ILI*, and *other*. Now we derive the posterior probability of *NI-ILI* on day *d* as the expected fraction of patients who have *NI-ILI* on that day:

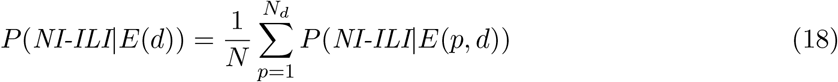

We derive *P*_*d*_(*influenza*|*E*(*d*)) and *P*_*d*_(*other*|*E*(*d*)) similarly. Finally, we assume that the prior probability of *NI-ILI* on day *d* + 1 is the posterior probability of *NI-ILI* on day *d*:

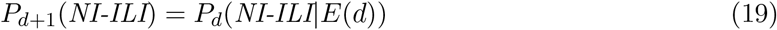

We can derive *P*_*d*+1_(*influenza*) and *P*_*d*+1_(*other*) similarly. Thus, starting with the baseline probabilities for *influenza*, *NI-ILI*, and *other* computed in the summer, we update them each day with the available evidence.

The probability of *NI-ILI* rises and falls with the probability of *influenza*. This is an artifact of the data: since *NI-ILI* and *influenza* have similar symptoms, *P* (*E*(*p, d*)|*NI-ILI*) will increase for patients with *influenza*, causing the likelihoods for *NI-ILI* to shadow the likelihoods for *influenza*. We will modify the computation of *NI-ILI* above with the goal of keeping the *NI-ILI* priors lower during an outbreak of *influenza* since we do not expect a simultaneous *NI-ILI* outbreak. Given the *NI-ILI* priors *P*_*d*_(*NI-ILI*) computed above, we will derive a new sequence of *NI-ILI* priors 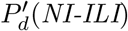. Set 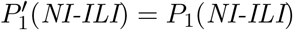 and assume *c* > 1. We run ODS for day *c* and derive 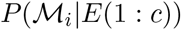 for each model 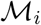 using the *NI-ILI* priors 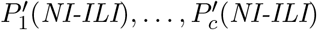. Let *P*_*c*_(*outbreak*|*E*(1: *c*)) be the probability that an outbreak is occurring on day *c* given *E*(1: *c*), and 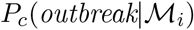 be the probability that an outbreak is occurring on day *c* given model 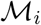 (as defined by Equation 8). We compute 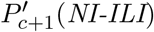 with:

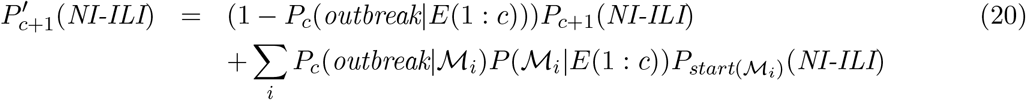

This equation says that we use the original *NI-ILI* prior on day *c*, *P*_*c*+1_(*NI-ILI*), weighted by the probability of no outbreak on day *c*, plus the sum over all models of the original *NI-ILI* prior on the start day of a model, *P*_*start*(*M*_**i**_)_(*NI-ILI*), weighted by the probability an outbreak is occurring on day *c* given model *M*_*i*_, *P*_*c*_(*outbreak*|*M*_*i*_) and the posterior of model *M*_*i*_ given the evidence through day *c*, *P*(*M*_*i*_|*E*(1: *c*)). The result of using Equation 20 is that when an influenza outbreak is occurring, the *NI-ILI* priors are weighted toward the values at the start of the most likely ongoing outbreak models, and do not have the behavior of increasing due to influence of the influenza outbreak.

## B Counts of Positive Influenza Lab Tests

## References

[1] Lipsitch M, Riley S, Cauchemez S, Ghani AC, Ferguson NM. Managing and Reducing Uncer-tainty in an Emerging Influenza Pandemic. New England Journal of Medicine. 2009;361(2):112–115.

[2] World Health Organization. Influenza Fact Sheet; 2003.

[3] Shaman J, Karspeck A. Forecasting Seasonal Outbreaks of Influenza. Proceedings of the National Academy of Sciences. 2012;109(50):20425–20430.

[4] Nsoesie E, Marathe M, Brownstein J. Forecasting Peaks of Seasonal Influenza Epidemics. PLoS Currents. 2013;.

[5] Ong JBS, Mark I, Chen C, Cook AR, Lee HC, Lee VJ, et al. Real-Time Epidemic Monitoring and Forecasting of H1N1-2009 Using Influenza-Like Illness from General Practice and Family Doctor Clinics in Singapore. PloS One. 2010;5(4).

[6] Wagner MM, Hogan WR, Chapman WW, Gesteland PH. Chief Complaints and ICD Codes. Handbook of Biosurveillance. 2006;.

[7] Ginsberg J, Mohebbi MH, Patel RS, Brammer L, Smolinski MS, Brilliant L. Detecting influenza epidemics using search engine query data. Nature. 2009;457(7232):1012–1014.

[8] Cooper GF, Villamarin R, Tsui FCR, Millett N, Espino JU, Wagner MM. A Method for Detecting and Characterizing Outbreaks of Infectious Disease from Clinical Reports. Journal of Biomedical Informatics. 2015;53:15–26.

[9] Wagner M, Tsui F, Cooper G, Espino J, Harkema H, Levander J, et al. Probabilistic, Decision-Theoretic Disease Surveillance and Control. Online Journal of Public Health Informatics. 2011;3(3).

[10] Tsui F, Dowling J, Vorhees R, Espino J, Wagner M. An Automated Influenza-Like Illness Reporting System Using Freetext Emergency Department Reports. ISDS. 2010;.

[11] Tsui F, Wagner M, Cooper G, Que J, Harkema H, Dowling J, et al. Probabilistic Case Detection for Disease Surveillance Using Data in Electronic Medical Records. Online Journal of Public Health Informatics. 2011;3(3).

[12] Provost F, Fawcett T. Data Science for Business: What You Need to Know of Data Mining and Data-Analytic Thinking. O’Reilly Media; 2013.

[13] Vynnycky E, White R. An Introduction to Infectious Disease Modelling. Oxford University Press; 2010.

[14] Carrat F, Vergu E, Ferguson NM, Lemaitre M, Cauchemez S, Leach S, et al. Time lines of Infection and Disease in Human Influenza: A Review of Volunteer Challenge Studies. American Journal of Epidemiology. 2008;167(7):775–785.

[15] Metzger KB, Hajat A, Crawford M, Mostashari F. How many Illnesses Does One Emergency Department Visit Represent? Using a Population-Based Telephone Survey to Estimate the Syndromic Multiplier. Morbidity and Mortality Weekly Report. 2004;53.

[16] for Disease Control C, Prevention. Flu Activity and Surveillance; 2016. Available from: www.cdc.gov/flu/weekly/fluactivitysurv.htm.

[17] Shaman J, Pitzer VE, Viboud C, Grenfell BT, Lipsitch M. Absolute Humidity and the Seasonal Onset of Influenza in the Continental United States. PLoS Biology. 2010;8(2).

[18] Gani R, Hughes H, Fleming D, Griffin T, Medlock J, Leach S. Potential Impact of Antiviral Drug Use During Influenza Pandemic. Emerging Infectious Diseases. 2005;11(9):1355–1362.

